# Apolipoprotein M-bound sphingosine-1-phosphate regulates blood-brain barrier paracellular permeability and transcytosis

**DOI:** 10.1101/684894

**Authors:** Mette Mathiesen Janiurek, Christina Christoffersen, Krzysztof Kucharz, Martin Lauritzen

## Abstract

The blood-brain barrier (BBB) is formed by the endothelial cells lining cerebral microvessels. Here, we report that the BBB permeability is modified by apolipoprotein M (apoM)-bound sphingosine 1-phosphate (S1P). We used two-photon microscopy to monitor changes in BBB permeability in apoM-deficient mice (apoM^−/−^), showing significant increases in paracellular BBB permeability to small molecules without structural changes in junctional complexes between endothelial cells. Lack of apoM-bound S1P increased vesicle-mediated transfer of albumin across endothelium of brain pial and penetrating arterioles, whereas transcytosis in capillaries and venules remained unchanged. S1PR1 agonist SEW2871 rapidly normalized BBB permeability along both the paracellular and transcellular routes in apoM^−/−^ mice. Thus, apoM-bound S1P maintains low paracellular BBB permeability for small molecules in all cerebral microvessels and low levels of adsorptive transcytosis in penetrating arterioles. Modulation of apoM/S1P-dependent signaling may be a novel strategy for the protection of brain endothelial cells to preserve the BBB function.

## INTRODUCTION

The blood-brain barrier (BBB) allows only certain molecules and cells to pass from the blood circulation to the brain. Molecules cross the BBB via the paracellular route by diffusion between endothelial cells (1), or by transcellular vesicular transport across the endothelium (2). The selectivity of the BBB towards small molecules is maintained by tight and adherens junctions between adjoining endothelial cells (1), whereas transport of large molecules is restricted by receptor-mediated interactions (2). Under normal conditions, the BBB maintains low permeability due to both the structural barrier maintained by junctional complexes and the low rate of transcytosis (2–5). In disease states, the loss of BBB function leads to increased permeability, which is viewed mainly as the consequence of disrupted junctional complexes at endothelial cell contact sites (6–8). However, recent evidence suggests an equal importance of transcytosis, as a failure in the homeostatic regulation of endothelial trans-cellular transport may aggravate brain pathologies (9–12).

Sphingosine-1-phosphate (S1P) is a bioactive sphingolipid synthesized by all mammalian cells, with blood-circulating S1P produced by vascular endothelial cells, platelets, and erythrocytes (13–16). S1P is a ligand for five membrane-bound G-protein-coupled receptors (S1PR1-5), and the brain endothelium expresses S1PR1-3 (16). The majority (~70%) of S1P in the blood circulation is transported by apolipoprotein M (apoM), a 26-kDa protein mainly associated with high-density lipoprotein (HDL) (17), while the remaining S1P fraction is carried by albumin (18, 19).

Depending on the carrier (i.e., apoM or albumin), S1P can preferentially activate S1PR1 or S1PR2 and S1PR3, which exert opposite effects (20). For example, S1PR1 stimulation inhibits neuroinflammation, whereas S1PR2 and S1PR3 stimulation activates pro-inflammatory signaling cascades (21–23). The onset and severity of sepsis and neuroinflammation (e.g., after brain trauma, ischemic stroke, or in brain cancer) correlate with reduced apoM-S1P plasma concentrations (24). Notably, these disorders are also associated with a loss of BBB integrity (6, 7, 10).

Lack of apoM impairs S1PR1 signaling and compromises the endothelial barrier in the lungs and brown adipose tissue (19, 25, 26), but the brain effects have remained unexplored. One study showed that conditional knock-out of S1PR1 increases the BBB permeability towards small molecules (27), but lack of S1PR1 expression is lethal in early stages of embryonic development and does not reflect apoM deficiency-mediated pathology well. Consequently, no data are available on how the BBB properties change with normal S1PR1 expression, but under conditions of reduced apoM-S1P levels that leads to S1PR1 hypostimulation.

Our objective was to determine the modulatory role of apoM for the BBB. We used two-photon microscopy to image the brains of apoM-deficient (apoM^−/−^) mice *in vivo* in order to characterize changes in BBB permeability in different categories of cerebral microvessels. Lack of apoM increased the BBB permeability to small molecules without affecting the ultrastructural components of junctional complexes and caused substantial increases in adsorptive transcytosis in pial and penetrating arterioles, but not in venules and capillaries. The increased permeability for both the paracellular and transcellular route was promptly reversed by systemic administration of SEW2871, a selective S1PR1 agonist. Thus, the loss of BBB function downstream of apoM-deficiency was mediated by S1PR1 hypostimulation. We suggest that modulation of the apoM-S1P signaling axis may be clinically useful for the development of protective strategies for the brain endothelium that may reduce secondary neural damage in both acute and chronic neurological conditions.

## MATERIALS AND METHODS

### Animals

All animal procedures were approved by the Danish National Ethics Committee and were in appliance with the ARRIVE guidelines. We used 14 to 20-week-old (22-28 g) female transgenic C57bl/6 apoM-deficient mice (apoM^−/−^) (19), and age- and gender-matched wild-type (WT) mice as the control group.

### Anesthesia

Anesthesia was induced by a single bolus intraperitoneal (i.p.) xylazine (10 mg/kg body weight [BW]) and ketamine (60 mg/kg BW) injection. During all steps of the surgical preparation, the anesthesia was maintained by i.p. injections of ketamine (30 mg/kg BW) at 25-minute intervals. Thirty minutes prior to imaging, the anesthesia was changed to continuous intravenous (i.v.) infusion of alpha-chloralose (50 μg/hour/g BW).

### Animal preparation for two-photon imaging

Animals were operated on as described previously (28). Briefly, a tracheotomy was performed for mechanical respiration (MiniVent Type 845, Harvard Apparatus) and to monitor exhaled CO_2_ (MicroCapstar End-tidal CO_2_ Analyzer, CWE). Two catheters were inserted: the first into the left femoral artery to monitor arterial blood pressure (Pressure Monitor BP-1, World Precision Instruments) and inject compounds, and the second into the vein for infusion of the anesthetic.

Next, a craniotomy was performed. The exposed skull was attached to a custom-made metal head bar with cyanoacrylate gel (Loctite Adhesives) and mounted onto the microscope imaging stage insert. The opening was made using a diamond dental drill (at 7000 rpm) over the somatosensory cortex (4 mm diameter; coordinates: 3 mm lateral, 0.5 mm posterior to bregma). During the procedure, the craniotomy and drill were cooled to room temperature repeatedly with artificial cerebrospinal fluid (aCSF; in mM: NaCl 120, KCl 2.8, Na_2_HPO_4_ 1, MgCl_2_ 0.876, NaHCO_3_ 22, CaCl_2_ 1.45, glucose 2.55, at 37°C, pH = 7.4) to prevent brain damage from excessive heating of the drill. The dura was removed and a drop of 0.75% agarose (type III-A, Sigma-Aldrich) applied onto the craniotomy. The opening was then secured with an imaging coverslip (Menzel-Gläser) and kept under aCSF to preserve humidity. To ensure physiological O_2_ and CO_2_ blood concentrations, prior to imaging, a 50 μl blood sample was collected via the arterial catheter (ABL blood gas analyzer, Radiometer) and the respiration rate and volume adjusted if necessary.

### Fluorescent probes

FITC-dextran (MW 10 kDa, 0.5%, 0.05 ml, Sigma-Aldrich), Alexa Fluor 488 (MW 0.643 kDa, 0.025%, 0.04 ml, Invitrogen), and sodium fluorescein (NaFluo, MW 0.365 kDa, 1%, 0.04 ml, Sigma-Aldrich) were each dissolved in sterile saline and administered via the arterial femoral catheter as single bolus injections. Bovine serum albumin (BSA) Alexa Fluor 488-conjugate (BSA-Alexa 488, Invitrogen) was administered via the arterial femoral catheter as a single bolus injection (0.05 ml 1%), together with TRITC-dextran (MW 65 kDa, 0.05 ml 1%,, Sigma-Aldrich) for delineation of the vasculature.

### Two-photon imaging setup

Two-photon imaging was performed using an SP5 upright laser scanning microscope (Leica Microsystems) equipped with a MaiTai Ti:Sapphire laser (Millennia Pro, Spectra Physics) using the 20× 1.0 NA water-immersion objective. The emitted light was split by an FITC/TRITC filter and collected by separate multi-alkali detectors after 525-560 nm and 560-625 nm band pass filters (Leica Microsystems).

### Monitoring paracellular permeability

The excitation wavelength for FITC-dextran, Alexa Fluor 488, and NaFluo was 850 nm (14 mWatt output power at the sample). Data were collected as 16-bit image hyperstacks at 512 × 512 pixel resolution (775 μm × 775 μm) in bidirectional scanning mode with a Z-step size of 5 μm (depth span = 114 μm) and 1-minute time interval between Z-stacks. The total recoding time for each fluorophore was 30 min. The recordings were exported to ImageJ (NIH, version 1.48u4) and Z-projected using maximal signal intensity projection. Regions of interest (ROIs) were chosen throughout the brain parenchyma at all locations deprived of vasculature for the measurement of fluorescent intensity. An increase in fluorescent intensity over the baseline (first projected Z-stack in time-lapse recording) in the parenchyma indicated crossing of the blood-circulating fluorophore into the brain.

### Monitoring transcellular transport

The animals were injected intravenously with BSA-Alexa 488. The excitation wavelength was 870 nm (17 mWatt output power at the sample). Data were collected as 16-bit hyperstacks at 2048 × 2048 pixel resolution (386.66 μm × 386.66 μm) in bidirectional scanning mode and triple frame averaging with a Z-step size of 2.50 μm, depth span 144 μm, and 7.5-minute interval between Z-stacks for a total recording time of 120 min. The data were exported to ImageJ and Z-projected using maximal signal intensity projection. Based on morphology and location, the vessels were classified as pial arterioles or venules, penetrating arterioles, post-capillary venules, ascending venules, or capillaries. To determine the vesicle surface density, the vessel surface area was calculated based on vessel diameter and length of vessel from which the BSA-Alexa 488 signal was counted.

### Transmission electron microscopy

The mice were anesthetized with bolus i.p. injections of xylazine (10 mg/kg BW) and ketamine (60 mg/kg BW), then perfused with Karnovsky fixation (2% glutaraldehyde and 4% paraformaldehyde). Brains were harvested and the cortex cut into 1 mm cubes and left for further fixation in 2% glutaraldehyde in 0.05 M phosphate buffer for 7 days. For transmission electron microscopy (TEM) staining, the samples were washed three times for 15-20 min in 0.12 M cacodylate buffer, followed by post-fixation in 1% osmium tetroxide (OsO_4_) and 0.05 M (1.5%) potassium ferricyanide (K_3_FeCn_6_) in 0.12 M cacodylate buffer for 1 hour at room temperature. Next, the samples were washed three times for 15 - 20 min in ddH_2_O and then dehydrated in an ethanol gradient (70%, 96%, and 3 × 15 min in absolute ethanol). Infiltration was performed with propylene oxide twice for 15 min, followed by incubation for 20 - 40 min in a resin epon/propylene oxide (Bie & Berntsen) gradient (1:3, 1:1, 3:1) and pure resin for 2 hours. Samples were embedded in resin at 60°C for 24 hours. Sections were cut at a thickness of 70 nm (Leica EM UC7), placed on copper grids (Leica), and stained with lead citrate (Leica Ultrostain II, Leica Microsystems) for 3 min and uranyl acetate (Leica Ultrostain I, Leica Microsystems) for 10 min using an automatic contrasting system (Leica EM AC20, Leica Microsystems). Imaging was performed using a CM100 transmission electron microscope (Phillips/FEI, ITEM software) at 9700× - 17500× and 46000-60000× magnification. Two samples from each mouse were imaged, and 20-30 capillaries from each sample were chosen randomly. ImageJ was used for the measurement of BBB morphology.

### Drug administration

SEW2871 (Sigma-Aldrich) was diluted in PBS (to 0.1 mg/ml in 1% DMSO) and administered as a single i.p. bolus injection (10 μg/g BW) 150 min prior to two-photon imaging or perfusion fixation.

### Statistical analysis

Statistical analysis was performed using Prism 7 (v7.0b, Graphpad) and R software (v3.5.1). The data were tested for normal distribution with Pearson’s normality test and either an unpaired two-tailed Student’s t-test with Welch’s correction (normally distributed data) or Mann-Whitney test (non-normally distributed data) was used. The TEM data underwent logarithmic transformation and was evaluated using one-way analysis of variance (ANOVA) followed by Tukey’s multiple comparison test. The sample size has been selected basing on our previous experiments using two-photon in vivo microscopy (28), in vivo electrophysiology (29) and TEM (19). The data consisted only from biological replicates, the number of independent biological replicates (N) is provided in results section and/or on figures. All analyses were performed in randomized order, with TEM analysis performed blinded to experimental conditions. No data has been excluded from analysis.

## RESULTS

### Deficiency in apoM^−/−^ increases BBB permeability

To assess the BBB with all structural constituents, including blood flow, we used two-photon fluorescence imaging *in vivo* in transgenic apoM-deficient mice (apoM^−/−^) mice (19). Following preparative surgery (Fig. 1a), the brain was imaged through an acute craniotomy over the somatosensory cortex (Fig. 1b).

**Figure 1:**
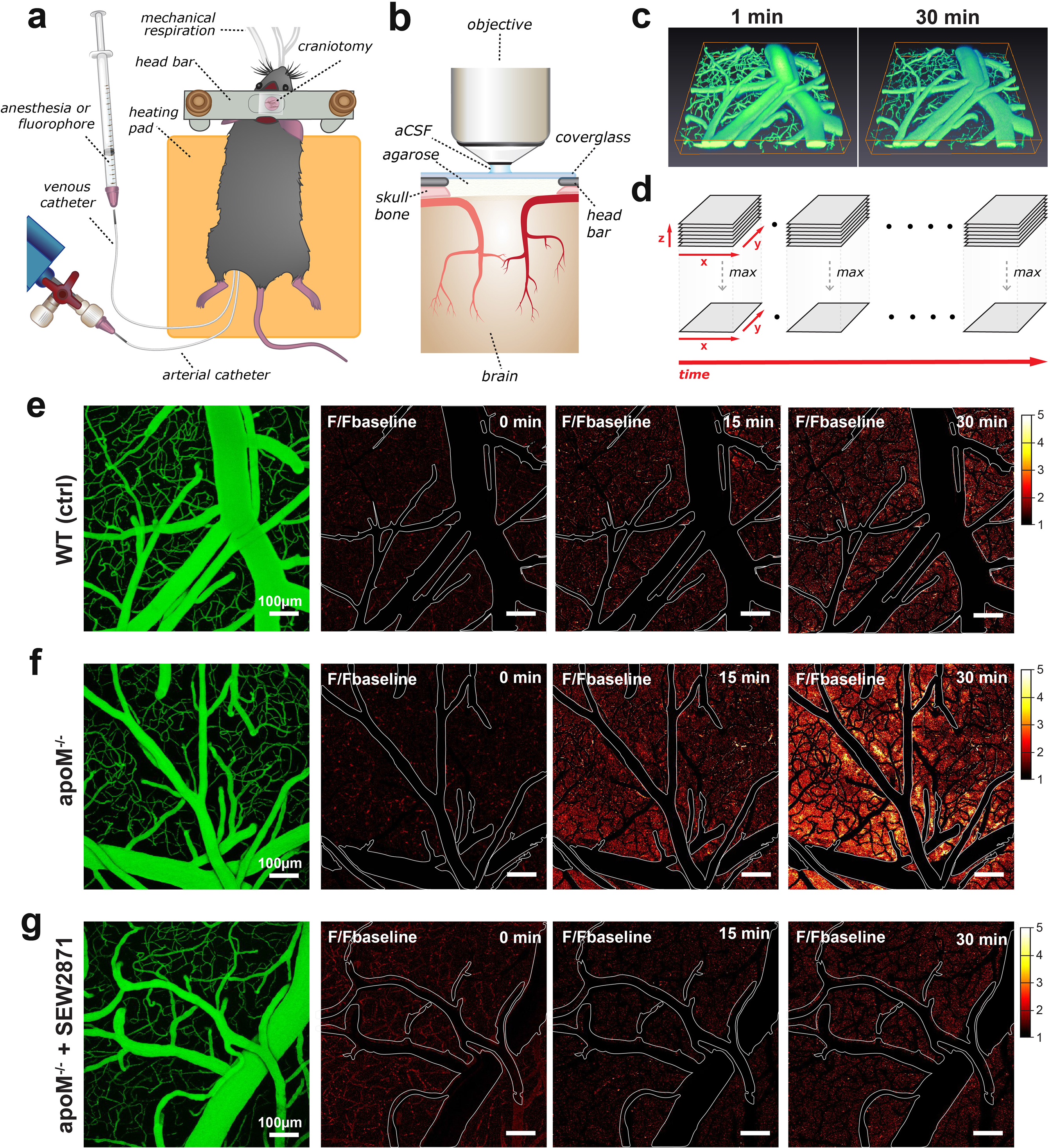
Two-photon imaging of paracellular permeability changes in apoM^−/−^ mice. **(a)** Schematic of a mouse after preparative microsurgery for two-photon imaging, with **(b)** craniotomy over the somatosensory cortex. **(c)** Three-dimensional reconstruction of the brain microvasculature from a single Z-stack after i.v. injection of NaFluo. **(d)** Each Z-stack corresponds to a single time point and underwent dimensionality reduction by maximum intensity projection along the Z-axis (depth). **(e-g)** Left, the general morphology of the vasculature visualized by blood-circulating NaFluo. The remaining panels show the relative fluorescence change over time in relation to the baseline (first projected Z-stack in recording). Contours delineate the main cerebral microvasculature. Compared to **(e)**WT mice, **(f)** the apoM^−/−^ animals had increased accumulation of NaFluo in the brain parenchyma over time, which was ameliorated by **(g)**treatment with S1PR1 agonist SEW2871.

The integrity of the BBB was assessed after an i.v. bolus injection of sodium fluorescein (NaFluo, 0.365 kDa). The fluorescence signal from blood-circulating fluorophore and brain parenchyma was simultaneously measured for 30 min in a series of Z-stacks of relatively large brain volume (750 μm × 750 μm × 114 μm) that contained pial and penetrating microvessels, including capillaries (Fig. 1c). Subsequently, each Z-stack underwent dimensionality reduction, i.e., maximum intensity projection along the Z (depth) axis (Fig. 1d).

The morphology of the vasculature delineated by blood-circulating NaFluo did not differ between WT and apoM^−/−^ mice (Fig. 1e-f). We detected progressive accumulation of NaFluo in the brain parenchyma in both WT and apoM-deficient mice. However, apoM^−/−^ mice exhibited considerably higher levels of fluorophore accumulation in the brain compared to WT mice already at 15 min after injection, with a clear difference at the 30-minute time-point (Fig. 1e-f, Video 1). This indicates that the absence of the apoM induces an increase in the paracellular permeability of the BBB.

### S1PR1 stimulation restores BBB permeability characteristics for small molecules

As apoM is the main carrier for S1P in the blood stream and S1P bound to apoM primarily activates S1PR1 on the endothelial cells (16, 20), we tested whether the increase in BBB permeability due to the lack of apoM was a consequence of the reduction of circulating S1P and S1PR1 hypostimulation. Animals were injected 150 min prior to imaging with a selective S1PR1 agonist, SEW2871. Compared to apoM^−/−^ mice, the SEW2871-treated animals did not exhibit increased permeation of NaFluo into the brain (Fig. 1g, Video 1), suggesting that the S1PR1 agonist rescued the abnormal apoM^−/−^ BBB phenotype. This result indicates that the apoM-dependent S1PR1 signaling deficiency is the primary cause underlying increased BBB permeability.

### Extent of BBB permeability induced by S1PR1 signaling deficiency

We next examined whether the BBB also exhibited increased permeability towards other molecules of different sizes and chemical properties. Each animal in the WT, apoM^−/−^, and apoM^−/−^ SEW2871-treated group was injected at 30-minute intervals with fluorophores of decreasing molecular size, starting with 10 kDa FITC-dextran, followed by 0.643 kDa Alexa Fluor 488 and 0.365 kDa NaFluo (Fig. 2a). After each injection, we imaged the same brain volume over time, with subsequent dimensionality reduction of the Z-stacks.

**Figure 2:**
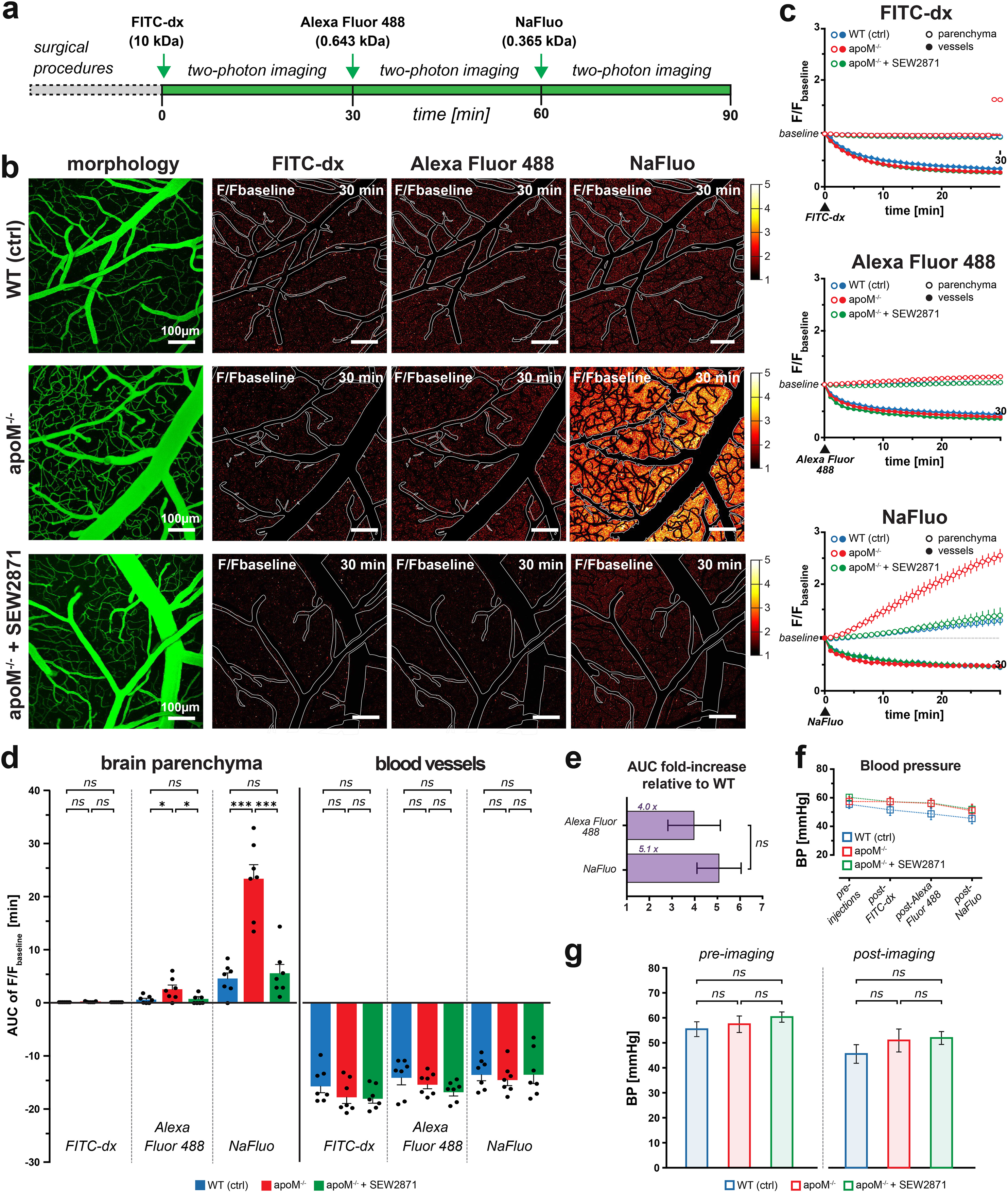
Extent of BBB permeability towards molecules of different sizes in apoM^−/−^ mice. **(a)** Experimental timeline. Bolus injections of fluorophore were separated by 30-minute time-lapse imaging intervals. **(b)** The vasculature morphology is outlined by a blood-circulating FITC-dx in the left column. The remaining panels show fluorescent molecule accumulation in the brain parenchyma for all tested fluorophores at 30 min post-injection. Imaging data are given as the relative change in fluorescence over time in relation to the baseline (first projected Z-stack in recording). **(c)** Changes in fluorescence intensity over time in brain parenchyma and the blood circulation expressed as a fraction of the baseline, which were used for **(d)** quantitative assessment of the total affect 30 min post-injection, i.e. area under the curve (AUC). Left bar plot, increased paracellular permeability of the BBB, i.e., the fluorescence increase in the brain parenchyma was evident for sodium fluorescein (0.365 kDa) and Alexa Fluor488 (0.643kDa) with no effect towards large molecule FITC-dextran (10 kDa). Right bar plot, no differences in the signal decrease in circulating fluorophores among all experimental groups. **(e)** The fold-increase in fluorescence intensity in the parenchyma in apoM^−/−^ mice compared to WT controls was similar between Alexa Fluor488 and sodium fluorescein. **(f)** The mean arterial blood pressure during imaging was **(g)** the same in all experimental groups. All data are presented as mean±SEM. *p<0.05, ***p<0.001.

To compare animal groups, we measured the changes in fluorescence signal over time from ROIs placed in the brain parenchyma and vasculature (n=7 mice in each group). For both tracer molecules, Alexa Fluor 488 and NaFluo, in all animal groups, we detected fluorescence increase over time in the parenchyma (Fig. 2b), and the effect was persistent without the signal reaching saturation (Fig. 2c). In contrast, there was no increase in brain fluorescence for FITC-dextran (Fig. 2b-c). To better characterize the fluorescence increase over time, we calculated the area under the curve (AUC) of the imaging traces, i.e., the cumulative effect of the fluorescence increase for duration of 30 min after fluorophore injection (Fig. 2d). We observed no differences in BBB permeability between WT and apoM^−/−^ mice for 10 kDa FITC-dextran (WT 0.0018±0.0014 min vs. apoM^−/−^ 0.066±0.035 min, p=0.1199), but the BBB in apoM^−/−^ mice was significantly more permeable to small molecules, such as Alexa Fluor 488 (WT 0.64±0.26 min vs. apoM^−/−^ 2.59±0.79 min, p<0.05) and NaFluo (WT 4.62±1.1 min vs. apoM^−/−^ 23.4±2.7 min, p<0.001) (Fig. 2d, Video 2-3). Noteworthy, the fold increase in BBB permeability in apoM^−/−^ mice over WT at baseline did not significantly differ between Alexa Fluor 488 and NaFluo (4.0 ±1.3 vs. 5.1±2.1, respectively, p=0.4709; Fig. 2e).

Treatment of the apoM^−/−^ mice with SEW2871 markedly reduced the leakage of both Alexa Fluor 488 and NaFluo compared to untreated apoM^−/−^ mice (0.59±0.30 vs. 2.59±0.79 min, p<0.05; and 23.4±2.7 min vs. 5.71±1.63 min, p<0.001; Fig. 2b-d, Video 3-4). The reversal of the dysfunctional BBB phenotype occurred relatively fast and at 150 min post-treatment with SEW2871, the BBB permeability was the same in apoM^−/−^ and WT mice (p=0.9130 and p=0.5914 for Alexa Fluor 488 and NaFluo, respectively; Fig. 2d).

Importantly, for all fluorophores and all investigated animal groups, the decrease in fluorescence signal in vessels (i.e., clearance of the fluorophore from the blood circulation) was the same during the whole imaging span (Fig. 2d). In addition, we found no differences in mean arterial blood pressure between animal groups before (pre-imaging) or after the administration of all fluorophores (post-imaging) (Fig. 2f-g). Thus, the increased fluorescence accumulation in the brains of apoM^−/−^ mice and reduced accumulation of fluorophores in the brains of SEW2871-treated animals were not caused by different kinetics of fluorophore clearance from the blood stream or differences in blood pressure.

These results show that apoM shortage causes S1PR1 signaling deficiency, increasing the BBB permeability towards small molecules (~0.3-0.7 kDa), and that the effect can be reversed by S1PR1 stimulation.

### Deficit in S1P/S1PR1 signaling does not alter tight junction ultrastructure

Increases in the BBB permeability towards small molecules may be indicative of defective structural elements that restrict paracellular diffusion across the BBB, i.e. morphological barriers, including junctional complexes (6–8). Therefore, we next assessed the ultrastructure of the BBB using TEM. We analyzed brain microvessels (< 6 μm) cross-sections from WT, apoM^−/−^, and SEW2871-treated apoM^−/−^ mice. In each vessel, we measured the thickness of the endothelium, basement membrane (both n=104 vessels; 4 mice), tight junctional complexes, and the endothelial cleft, the percentage coverage of endothelial cell contact sites by junctional complexes (all n=164, 4 mice), and the endothelial cell area (n=104 vessels, 4 mice) (Fig. 3, Fig. S1). The width, length, and coverage of junctional complexes were the same for all three groups of mice (Fig. 3a-b, Fig. S1a), suggesting that the compromised BBB was not caused by a structural defect, and that the rescue effect of SEW2871 was not mediated by alteration of the morphology of the BBB structural components. However, we detected an ~10% increase in the thickness of the endothelium in S1P-signaling-deficient mice compared to controls (apoM^−/−^ 0.078±0.0026 μm vs. WT 0.071±0.0025 μm; p<0.05) and the rescue group (apoM^−/−^+SEW2871 0.070±0.0025 μm; p<0.05; Fig. 3c). Notably, the stimulation of S1PR1 in apoM^−/−^ mice decreased the endothelial cell area (5.80±0.37 vs. 4.24±0.23 μm^2^; p<0.001), with a strong trend for the cell area in apoM^−/−^ being larger than WT (5.80±0.37 vs. 4.78±0.24 μm^2^; p=0.068; Fig. 3d). These changes were relatively small, and they are unlikely to underlie the increased BBB permeability for small molecules. Altogether, our results suggest that the increased permeability observed in apoM^−/−^ mice is not mediated by ultrastructural changes at the level of the junctional complexes.

**Figure 3:**
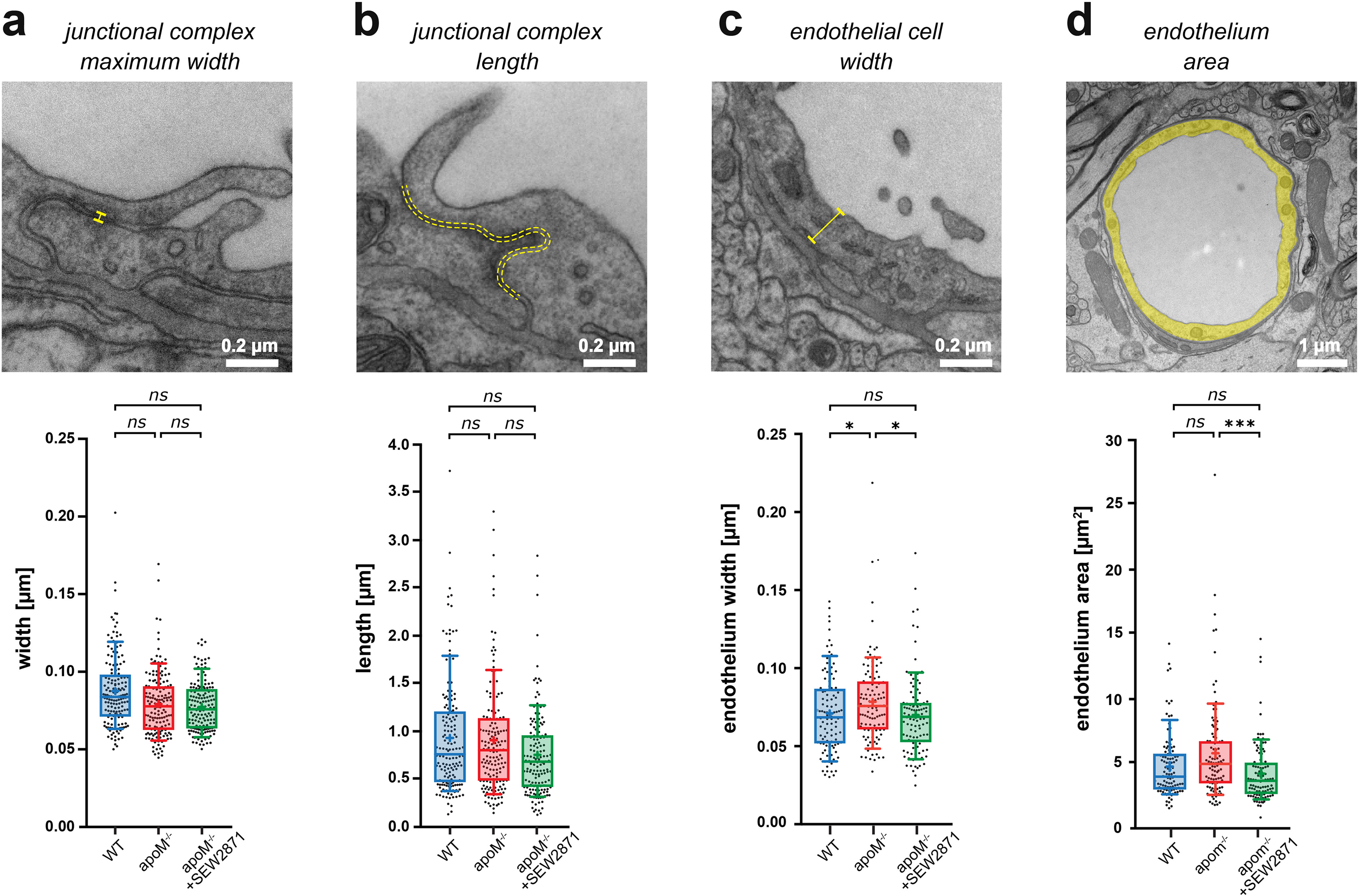
Small changes in endothelium cell morphology, but not junctional complexes, accompany increased BBB paracellular permeability in apoM^−/−^ mice. TEM ultrastructure assessment of the structural components of the BBB revealed no changes in the **(a)** width or **(b)** length of junctional complexes, with **(c)** a small increase in endothelial thickness in apoM^−/−^ mice compared to WT mice and a decrease in SEW2871-treated apoM^−/−^ mice. **(d)** A small decrease in endothelial cell area was observed upon SEW2871 treatment in apoM^−/−^ mice. Data are presented as interquartile distributions with median (horizontal line) and mean (plus sign). The box whiskers indicate 10-90% percentiles. *p<0.05, ***p<0.001.

### ApoM / S1PR1 signaling deficit increases transcellular transport

Next, we examined whether lack of apoM leads to increased transcytosis in the BBB. The animals were injected with BSA-Alexa 488 to monitor vesicular transport across the endothelium (9) and 65 kDa TRITC-dx to delineate the vessel lumen. Following the injections, the animals were continuously imaged for 120 min in hyperstack mode with subsequent dimensionality reduction (as in Fig. 1d). In all animal groups, and as early as 30 min post-injection, we detected BSA-Alexa 488-containing vesicles at the BBB interface (Fig. 4). Compared to WT animals, apoM^−/−^ mice exhibited increased albumin in endothelial cells, indicating that vesicular transport was increased in the absence of apoM. Furthermore, the treatment of apoM^−/−^ animals with SEW2871 reversed the observed effect, suggesting that the same apoM-dependent mechanism underlies the previously observed increase in permeability for small fluorescent tracers, as well as elevated transcytosis. Noteworthy, in apoM^−/−^ mice, we observed an increase in the presence of cells containing vesicular albumin in the brain parenchyma (Fig 4). The size of the cell bodies corresponded to activated (phagocytic) microglia (30), and their presence was consistent with ongoing neuroinflammation during S1PR1 hypostimulation (21–23).

**Figure 4:**
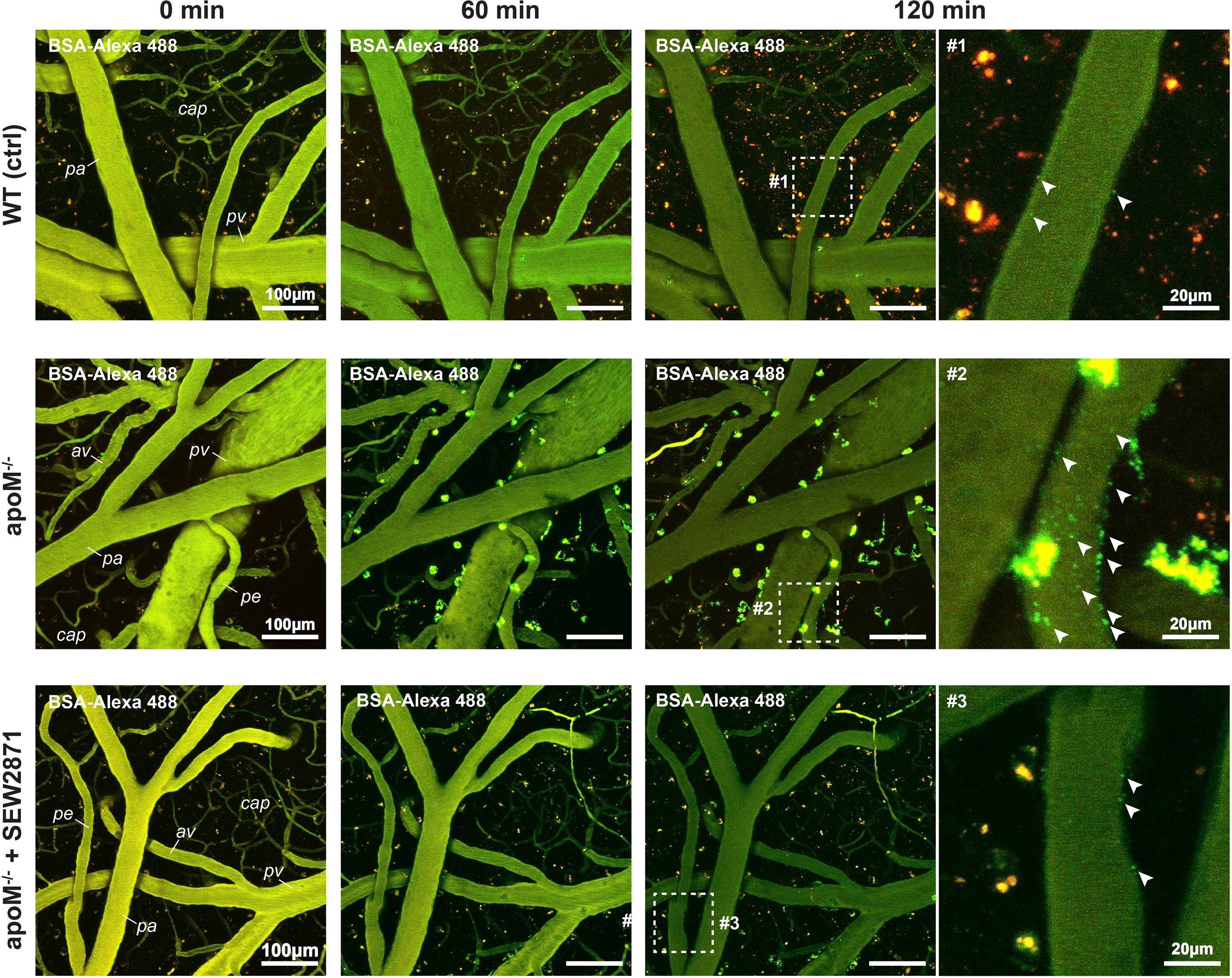
Increase in BSA-Alexa 488 uptake indicates an increase in transcellular transport in apoM^−/−^ mice. Two-photon assessment of albumin uptake (green puncta) 60 and 120 min post i.v. co-injection with BSA-Alexa 488 and 65 kDa TRITC-dextran. Arrowheads indicate numerous puncta of BSA-Alexa 488-positive vesicles at the BBB interface in apoM^−/−^ mice but only sparse labeling in WT and SEW2871-treated animals. pa=pial arteriole; pe=penetrating arteriole; cap=capillaries; pv=pial venule; av=ascending venule.

### S1P signaling deficit elevates transcytosis only in arterioles

Next, we assessed whether the increase in transcellular transport caused by the S1PR1-signaling deficiency was the same for all vessel types. The number of vesicles was quantified over time for each individual vessel type: pial arterioles, penetrating arterioles, capillaries, post-capillary venules, ascending venules, and pial venules (n = 5 mice). The data are expressed as the vesicle surface density, i.e., the number of vesicles per square micrometer of vessel surface area (Fig. 5a). In all experimental groups, the distribution of BSA-Alexa 488-positive vesicles was dependent on vessel type and diameter. At 120 min post-injection, the highest vesicle density in WT mice was observed in capillaries and gradually decreased with vessel diameter, regardless of the vessel type (i.e., arterioles or venules) (Fig 5a). In apoM^−/−^ mice, the deficiency in S1PR1 signaling selectively increased transcytosis in pial and penetrating arterioles, but not in venules and capillaries. The effect was most pronounced in penetrating arterioles, where the vascular vesicle density in apoM^−/−^ mice was 10-fold higher than in WT mice (apoM^−/−^ 0.30±0.071 vs. WT 0.038±0.009 vesicles/100 μm^2^; p<0.01), and three times higher than for pial arterioles (apoM^−/−^ 0.099±0.0023 vs. WT 0.035±0.0011 vesicles/100 μm^2^; p<0.05; Fig. 5b). The level of transcytosis did not change in apoM^−/−^ mice in the capillaries, venules, or pial veins (p=0.2306, p=0.3562, and p=0.9546, respectively).

**Figure 5:**
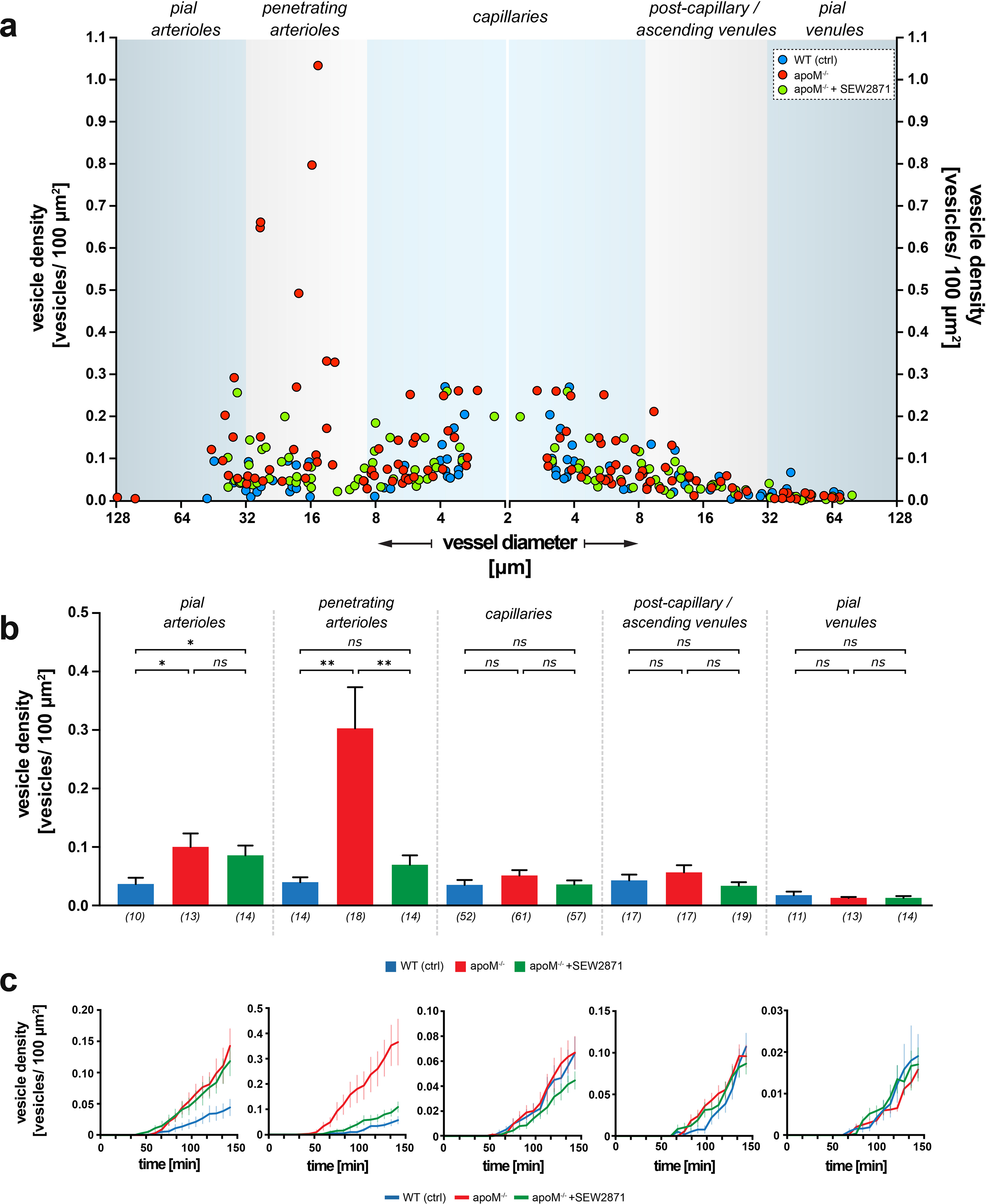
Heterogeneous susceptibility to increased transcytosis in apoM^−/−^ mice. **(a)** Scatter plot showing the distribution of the surface vesicle densities of individual vessels plotted against respective vessel diameters 120 min post BSA-Alexa 488 injection. The blue areas represent the morphological division into distinct vessel types. Data from capillaries were mirrored to preserve the order of vessel diameter change in the continuous vascular tree. **(b)** Quantitative analysis showed the increase in albumin uptake in apoM^−/−^ mice that is present only in the arterial part of the vascular tree and absent in capillaries and venules. Only penetrating arterioles responded to SEW2871 treatment. Numbers in brackets are the number of vessels across five mice in each group. The data are presented as mean±SEM. **(c)** Average traces showing the kinetics of the vesicle density change for WT, apoM^−/−^, and apoM^−/−^ SEW2871-treated mice with respect to different vessel types. The data are presented as mean±SEM. *p<0.05, **p<0.01.

### Penetrating, but not pial arterioles respond to SEW 2871 treatment

S1PR1 stimulation with SEW2871 led to normalization of the transcellular transport of albumin in the penetrating arterioles of apoM^−/−^ mice to the level of WT controls (apoM^−/−^+SEW2871 0.68±0.016 vs. WT 0.038±0.009 vesicles/100 μm^2^, p=0.1156; Fig. 5b). In contrast, SEW2871 treatment did not decrease transcytosis in pial arterioles (apoM^−/−^+SEW2871 0.84±0.017 vs. apoM^−/−^ 0.99±0.023 vesicles/100 μm^2^; p=0.4879), and albumin transport was still significantly elevated compared to WT mice (apoM^−/−^+SEW2871 0.84±0.017 vs. WT 0.35±0.011 vesicles/100 μm^2^; p<0.05). Lastly, we assessed the kinetics of the vesicular density increase over time with respect to animal group and vessel type. In all vessel types and animal experimental groups, the increase in BSA-Alexa 488 vesicle surface density was persistent and did not reach saturation at 120 min, indicating that the results were not obscured by overloading the transcellular transport with BSA-Alexa 488 (Fig. 5c).

These findings show that transcellular transport is upregulated in apoM-deficient mice, and that there is not only heterogeneity in the vasculature susceptibility towards increased permeability, but also towards which vessels can be rescued by stimulation of the apoM-dependent S1PR1 signaling pathway.

## DISCUSSION

Understanding the influence of blood-borne signaling molecules on brain endothelial cells is critical for an understanding the BBB function. We report that lack apoM-bound S1P increased the BBB permeability for small molecules without changes in size, structure, magnitude, or subcellular localization of tight junction complexes. Furthermore, we report an up to 10–fold increase in adsorptive transcytosis in the endothelial cells of pial and penetrating arterioles, but not in venules and capillaries. Thus, ApoM modulates the flux of small and large molecules to different extents in different vascular segments. Finally, the increased BBB permeability for small molecules and transcytosis-mediated large proteins could be rescued by selective stimulation of the S1PR1.

From among the exogenous tracers typically used in BBB paracellular permeability studies, we chose three molecules with different sizes, NaFluo and Alexa Fluor 488 that readily diffuse via the BBB, and FITC-dextran, which is reported to have low BBB permeability (31, 32). The loss of BBB integrity in apoM^−/−^ mice was observed for small tracer molecules <0.7 kDa, but no significant leak was observed for 10 kDa dextran, which is in accordance with previous findings in S1PR1–knock-out mice (27). However, in contrast to Yanagida and colleagues (27), we show that it is not the complete absence of brain endothelial S1PR1 signaling, but a lack of a specific fraction of S1P circulating in the blood (i.e., apoM-bound S1P) that induces a significant increase in BBB permeability. This occurred despite the presence of active S1PR1 and the remaining S1P circulating in the blood bound to albumin or in an unassociated form in apoM^−/−^ mice (19).

S1PR1 is implicated in the maintenance of BBB integrity by stabilization of the actin cytoskeleton that anchors junctional complexes at the endothelial membrane cell contact sites (15, 33). The increased permeability along the paracellular pathway occurs most commonly in response to severe damage (34–36) and is typically reported to be secondary to the ultrastructural abnormalities of junctional complexes, such as truncated morphology during acute BBB breakdown, e.g. in stroke, sepsis, and brain tumors (6, 7, 9), and in diabetes (8). While S1PR1–knock-out abolishes the normal structure of tight junctions and the composition of proteins (27), we show that the increases in BBB permeability caused by apoM-S1P signaling deficiency can occur without reduced cytoskeletal association or alteration of the barrier ultrastructure. This finding is in agreement with another study that reported a lack of ultrastructural changes in lung endothelium in the absence of apoM, whereas the cellular barrier function was impaired (37). Notably, a significant increase in paracellular permeability without apparent changes in junctional complex morphology has been observed by others following osmotic shock (34) and in claudin-5-deficient mice (38). The loss of BBB integrity in apoM^−/−^ mice appeared to be to a lesser extent than in acute pathology (e.g., traumatic brain injury or ischemic stroke), as the BBB permeability towards 10 kDa dextran did not increase in apoM^−/−^ mice (10, 39). Yet, even with is morphology intact, the BBB with a persistent increase in permeability may still contribute to brain pathology (40).

Though paracellular permeability restriction is typically viewed as the main functional constituent of BBB function, the low prevalence of transcytosis is equally important to preservation of the brain microenvironment (9–12). Recently, Mfsd2a was identified as a critical protein in the suppression of transcytosis. Similar to apoM^−/−^, a lack of Mfsd2a leads to an increase in vesicular transport across the endothelium that compromises the BBB without structurally opening the paracellular route (41) or changing junctional protein composition (42). Furthermore, the increase in BBB permeability can occur via both the transcellular and paracellular routes, but at different time points; the increase in transcytosis rate precedes paracellular leakage in stroke (9), and selective dysregulation of the transcellular pathway can occur while the tight junctions are preserved (12). Thus, the intact morphology of junctional complexes, as shown here, is not necessarily indicative of preserved BBB properties. Noteworthy, the lack of apoM-S1P lead to an increased rate of transcytosis, though only in pial and penetrating arterioles, which could be explained by non-homogeneous expression of S1PR1, which is higher in arteriole endothelial cells than the endothelium of capillaries and venules (43).

Different G protein-coupled receptors are activated downstream of S1PR1 and S1PR2 (44, 45), which results in opposite regulatory outcomes, such as in neuroinflammation (21–23). The S1P bound to ApoM primarily activates S1PR1, which underlies the preservation of endothelial barrier integrity in the kidneys and lungs (19, 25). Conversely, activation of S1PR2 has been implicated as a major factor in the induction of vascular permeability following stroke (46). In a healthy brain, regardless of the carrier, S1PR1 and S1PR2 are constantly stimulated by blood-circulating S1P, which is suggested to underlie the homeostatic regulation of endothelial integrity (15, 16). We show that reduction of apoM upstream of S1PR1 activation is a clinically relevant mechanism that may shift the balance towards S1PR2 activation, resulting in increased BBB permeability. Consequently, the observed impairment of the BBB cannot simply be linked to a reduction in the amount of circulating S1P, but more specifically to the lack of apoM-bound S1P in favor of albumin-bound S1P in the blood. This notion is further supported by the result of the selective stimulation of S1PR1, which restored the BBB permeability in apoM^−/−^ mice to the level of healthy controls.

In summary, ApoM-associated S1P is critical for maintaining low paracellular and transcellular permeability at the BBB. The pronounced increase in transcytosis in apoM^−/−^ mice stresses the importance of considering the development of protective strategies for this pathway in order to regulate the immediate microenvironment of brain cells and reduce secondary neural damage in both acute and chronic neurological conditions.

## Acknowledgements

We thank the Core Facility for Integrated Microscopy at University of Copenhagen, especially Klaus Qvortrup, for sample preparation for TEM. We thank Micael Lønstrup from the Department of Neuroscience at University of Copenhagen for technical guidance in the animal surgeries. This work was supported by the Lundbeck Foundation, the Danish Council for Independent Research (Medical Sciences), and a Nordea Foundation Grant to the Center for Healthy Aging.

## SUPPLEMENTARY FIGURE LEGENDS

**Supplementary Figure 1:**
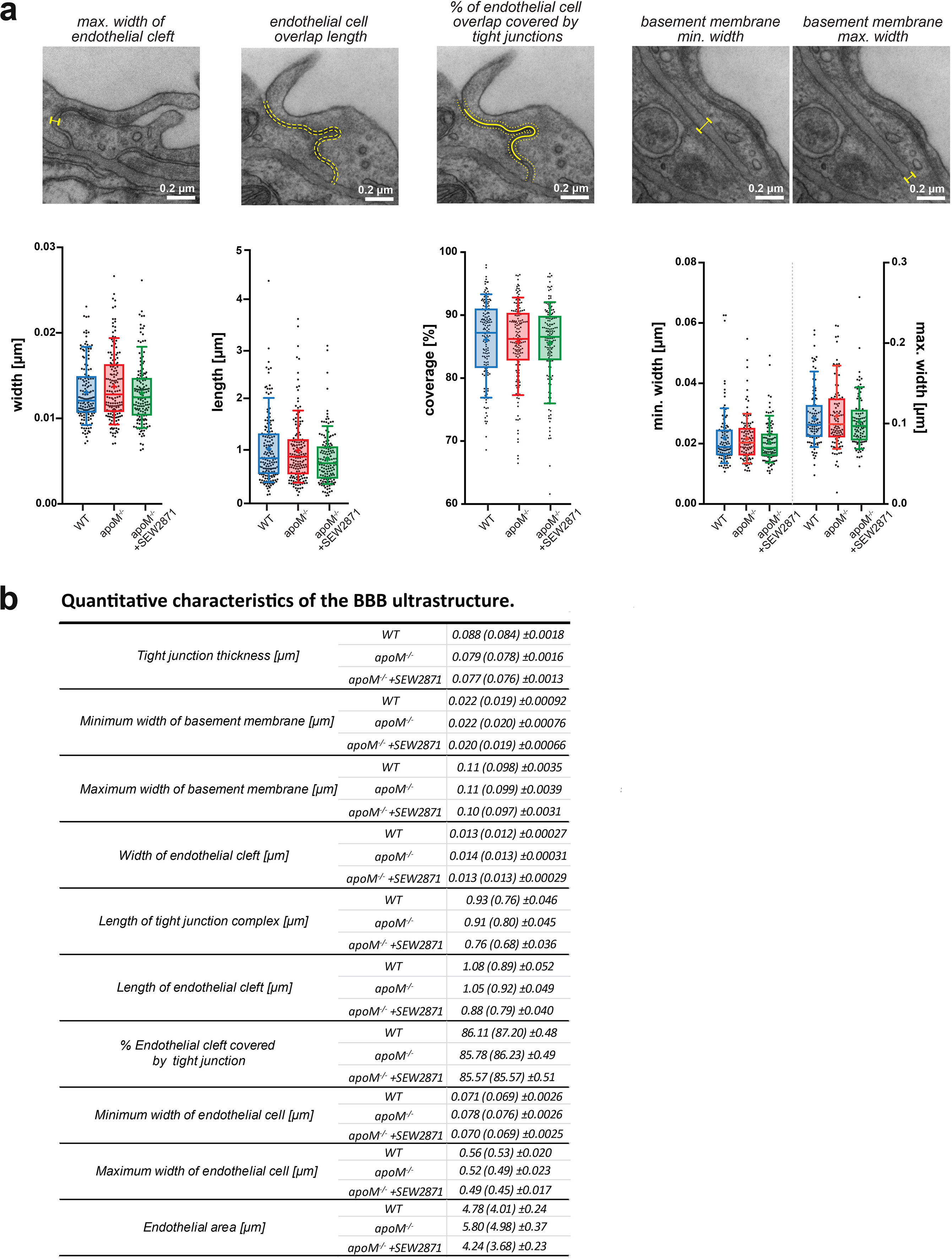
Complementary BBB measurements by TEM. **(a)** Neither ApoM-S1P deficit nor treatment with S1PR1 agonist SEW2871 affected the width of junctional complexes and length of the endothelial cleft, the proportion of endothelial cleft covered by junctional complexes (%), or the minimum and maximum thickness of the basement membrane. Data are presented as interquartile distributions with median (horizontal line) and mean (plus sign). The box whiskers indicate 10-90% percentiles. **(b)** Summary table of all TEM micrograph measurements. Data are presented as mean with (median) ± SEM.

## SUPPLEMENTARY VIDEOS LEGENDS

**Video 1: Lack of apoM leads to a reversible increase in BBB paracellular permeability.** Accumulation of NaFluo in brain parenchyma presented as fluorescence increase relative to the baseline. Fluorophore injections occurred 1 minute before the first recorded imaging frame.

**Video 2: WT mice BBB paracellular permeability towards NaFluo, Alexa488, and 10kDa FITC-dx.** Accumulation of fluorophores in brain parenchyma presented as fluorescence increase relative to the baseline. For each time-lapse recording, the respective fluorophore was injected 1 minute before the first recorded imaging frame.

**Video 3: apoM^−/−^ mice BBB paracellular permeability increase towards NaFluo, Alexa488 but not 10kDa FITC-dx.** Accumulation of fluorophores in brain parenchyma presented as fluorescence increase relative to the baseline. For each time-lapse recording, the respective fluorophore was injected 1 minute before the first recorded imaging frame.

**Video 4: apoM^−/−^ SEW2781-treated mice exhibit normal BBB paracellular permeability towards NaFluo, Alexa488 and 10kDa FITC-dx.** Accumulation of fluorophores in brain parenchyma presented as fluorescence increase relative to the baseline. For each time-lapse recording, the respective fluorophore was injected 1 minute before the first recorded imaging frame.

